# RNAElectra: An ELECTRA-Style RNA Foundation Model for RNA Regulatory Inference

**DOI:** 10.64898/2026.03.15.711950

**Authors:** Ke Ding, Lixinyu Liu, Brian Parker, Jiayu Wen

**Affiliations:** John Curtin School of Medical Research, Australian National University, Canberra, Australia; The Shine-Dalgarno Centre for RNA Innovation, Australian National University, Canberra, Australia; ARC Centre of Excellence for the Mathematical Analysis of Cellular Systems, Canberra, Australia; School of Computing, Australian National University, Canberra, Australia; Biological Data Science Institute, Australian National University, Canberra, Australia

**Keywords:** RNA regulation, RNA foundation model, genomic large language models, replaced-token detection, single-nucleotide resolution, RNA structure, RNA modifications, RNA–protein interactions, interpretability

## Abstract

**Background:** RNA regulation programs gene expression through sequence-encoded mechanisms—including RNA structure formation, protein binding, chemical modification, and RNA–RNA targeting—whose rules span nucleotide-scale motifs and longer-range context. RNA foundation models aim to learn transferable representations from large RNA corpora, but most rely on masked language modeling (MLM), where the model loss is computed on a small subset of positions and training relies on artificially corrupted inputs that are absent at downstream inference, introducing a pretraining–downstream discrepancy.

**Results:** We introduce RNAElectra, a single-nucleotide–resolution RNA foundation model pretrained on diverse non-coding RNAs from RNAcentral using ELECTRA-style replaced-token detection (RTD). RTD trains a discriminator with a loss defined over all input positions on realistically corrupted sequences, providing dense supervision that better aligns pretraining with sequence-to-function fine-tuning. RNAElectra combines nucleotide-resolution tokenization with an efficient attention design to capture local regulatory motifs and longer-range dependencies within a single reusable backbone. Using a unified, sequence-only fine-tuning pipeline without task-specific architectures or auxiliary inputs, RNAElectra demonstrates strong cross-task generalization across benchmarks and downstream evaluations spanning RNA structure and function, RNA–protein and RNA–RNA interactions, RNA modification landscapes, and quantitative regulatory readouts such as translation efficiency and mRNA stability, outperforming widely used RNA foundation model baselines on the majority of evaluated tasks. In addition to predictive performance, RNAElectra supports interpretability by enabling analysis of learned representations and sequence determinants underlying model predictions.

**Conclusion:** RNAElectra establishes RTD pretraining as a practical alternative to MLM for RNA foundation modeling, coupling dense position-wise supervision with an efficient architecture to deliver broadly transferable RNA representations. This framework provides a reusable backbone for RNA regulatory prediction and supports sequence-level RNA engineering and design.

## 1 Background

RNA is a central layer of biological regulation: it carries coding information for protein synthesis while also encoding regulatory signals that shape RNA processing, stability, localization, and translation [1–3]. A distinctive feature of RNA regulation is that it is written in a compact, multi-scale “regulatory grammar”: short nucleotide motifs recruit proteins and guide RNAs, chemical modifications tune recognition and decay, and long-range dependencies emerge through folding and context, collectively determining functional outcomes for each transcript [1–3]. Beyond its endogenous roles, RNA has increasingly become a major focus in medicine and biotechnology, acting as both a molecular readout of disease states and a programmable substrate for intervention [4, 5]. RNA-based modalities such as mRNA therapeutics and vaccines, antisense oligonucleotides, and guide RNAs rely on sequence-encoded determinants to achieve robust activity [4–7]. Yet RNA regulation spans multiple RNA classes and regulatory layers of control, from pre-mRNA splicing to post-transcriptional control, creating a high-dimensional space of context-dependent sequence rules [7, 8]. This combination of practical demand and biological complexity motivates computational approaches that can learn generalizable sequence-to-function principles and transfer them across diverse regulatory settings [9–11].

Large language models (LLMs) have established self-supervised pretraining as a general paradigm for learning reusable representations from discrete sequences, and these ideas transfer naturally from natural language to biological polymers [12]. RNA foundation models take RNA sequence as input and aim to capture reusable features that support diverse downstream predictions of RNA structure and regulation [12]. Early work such as RNA-FM showed that Transformer-based pretraining on large RNA collections yields informative features for RNA structure and function prediction [10]. Subsequent models increased scale and broadened training data sources (e.g., RiNALMo and AIDO.RNA) targeting broad task transfer via pretraining on tens of millions of RNA sequences [13, 14]. Other approaches incorporate stronger structural and evolutionary priors: RNABERT [15] and RNAFormer [16] integrate structure-aligned objectives and 2D latent representations; RNA-MSM [17] leverages multiple sequence alignments to encode covariation; and RNAErnie [18] introduces biologically motivated, motif-aware masking strategies. Complementary to these “global” models, region-specific language models focus on well-defined regulatory contexts, including untranslated regions (i.e., UTR-BERT [19], UTR-LM [20]) and coding or splicing-centric semantics (CodonBERT [21] and SpliceBERT [22]). More recent efforts explore long-context scalability and RNA sequence generation for design (HydraRNA [23], RNAGenesis [24]), and unified modeling across interconnected regulatory layers (LAMAR [25]).

Despite rapid progress, several limitations still constrain practical utility for RNA regulatory inference. First, most RNA foundation models rely on masked language modeling (MLM), where the loss is computed on a small subset of masked positions and training operates on artificially corrupted inputs that are absent at downstream inference. This mismatch between the pretraining signal and downstream usage can reduce the effectiveness of position-wise learning, particularly for RNA mechanisms whose signals are subtle and distributed across many positions (e.g., motif cores together with informative flanking context and structure-dependent constraints) rather than a small set of isolated positions. Second, many existing models tokenize RNA into *k*-mers or longer segments to improve efficiency, but this can blur single-nucleotide effects that are central to regulatory interpretation, variant impact analysis, and rational sequence editing. This is particularly relevant for motif-driven regulation, where information content varies substantially by base position and single-nucleotide substitutions can have disproportionate functional consequences. In addition, downstream pipelines are often heterogeneous across tasks, with task-specific heads, auxiliary features, or preprocessing choices that reduce portability and reuse across datasets and experimental settings, and can complicate attribution of gains to the pretrained representation itself.

Here we propose RNAElectra, an RNA foundation model that addresses these limitations by adopting ELECTRA-style replaced-token detection (RTD) for RNA pretraining. Motivated by our prior ELECTRA-style genomic foundation modeling work for DNA (NucEL) [26], we adapt RTD to RNA by pretraining RNAElectra from scratch on RNA sequences to better align pretraining with downstream sequence-to-function prediction. In this generator–discriminator scheme, a lightweight generator proposes plausible nucleotide replacements at selected positions, and a discriminator is trained to predict at every position whether the token is original or has been replaced. By defining the loss over all input positions on realistically corrupted sequences, RTD provides dense supervision that better matches downstream inference on fully observed (unmasked) sequences, where predictions often depend on distributed cues across many nucleotides. RNAElectra operates at single-nucleotide resolution and uses an efficient attention design that integrates local and global context modeling to capture both short-range regulatory motifs and longer-range dependencies within a single backbone. RNAElectra is trained and applied with a unified, sequence-only pipeline without task-specific architectural modifications or auxiliary inputs, simplifying transfer across heterogeneous tasks and datasets. Concretely, we pretrain RNAElectra on diverse non-coding RNAs from RNAcentral [27] and benchmark it across broad evaluations spanning RNA structure, binding and modification signals, and quantitative regulatory readouts such as translation efficiency and mRNA stability, where it outperforms widely used RNA foundation model baselines on the majority of tasks. We further complement predictive benchmarking with embedding-space analyses and motif discovery on RNA-protein binding predictions to probe the sequence determinants captured by the model.

## 2 Results

### 2.1 RNAElectra: efficient ELECTRA pretraining learns RNA’s regulatory grammar at single-nucleotide resolution

RNAElectra is an ELECTRA-style RNA foundation model pretrained with replaced-token detection (RTD), in which a lightweight generator proposes plausible nucleotide substitutions and a discriminator learns to detect replaced tokens at each position (Fig. 1a). The model operates at single-nucleotide resolution and uses global attention in every layer, enabling sensitivity to motif-scale signals while retaining long-range context integration. We pretrained RNAElectra on roughly 44 million curated RNA sequences from RNAcentral, an integrated resource that aggregates non-coding RNA sequences and annotations from many specialist databases across diverse species (Fig. 1b), totaling ∼20 billion tokens. We then fine-tuned the same backbone under a unified, sequence-only protocol across downstream tasks (Fig. 1c).

**Fig. 1.**
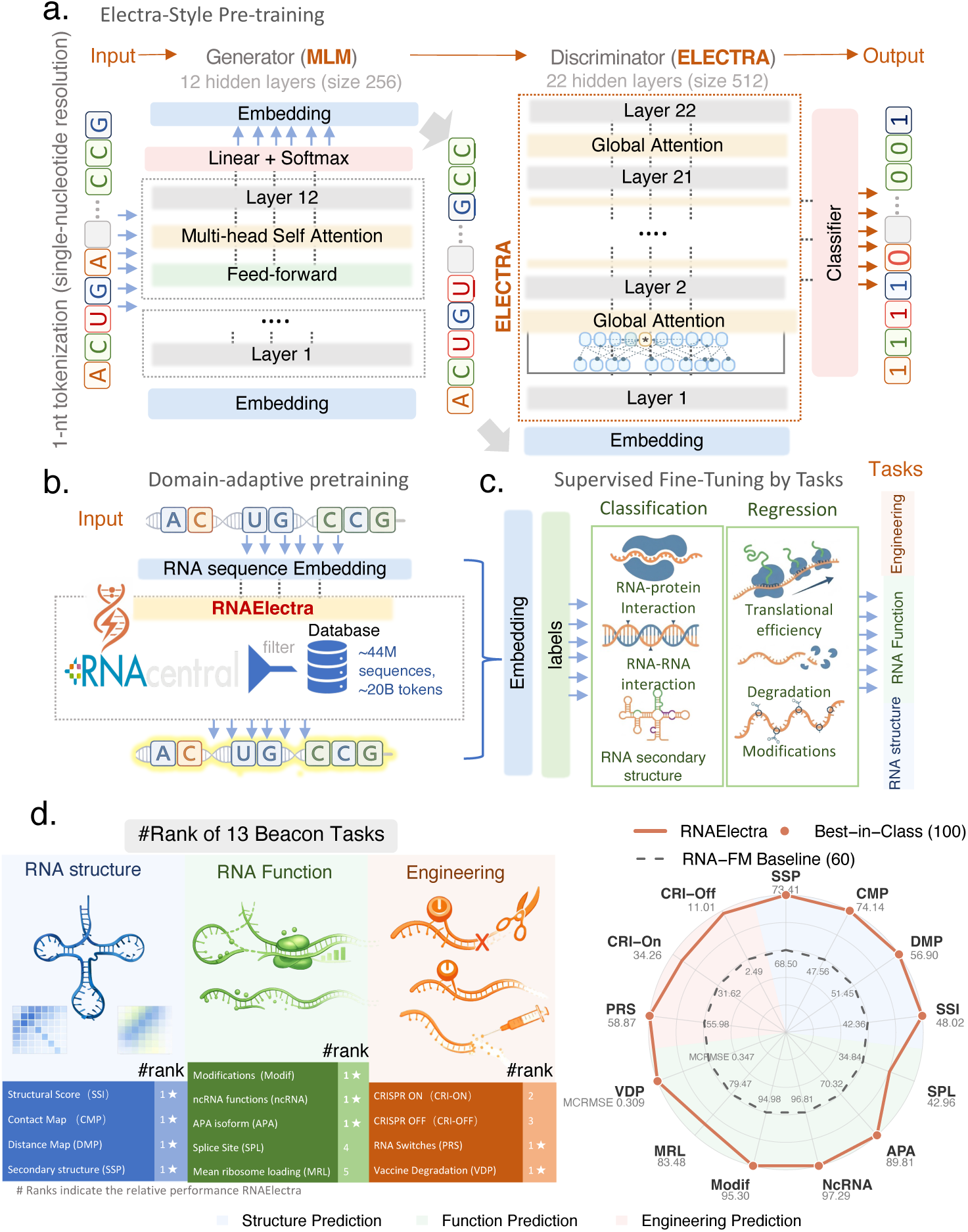
RNAElectra overview: architecture, pretraining, and benchmark performance. **(a)** ELECTRA-style pretraining for RNA at single-nucleotide resolution. A lightweight masked language model (MLM) *generator* (12 layers, hidden size 256) proposes context-dependent nucleotide replacements, while a deeper *discriminator* (22 layers, hidden size 512) is trained with replaced-token detection to classify, at each position, whether the observed nucleotide is original or replaced. Both the generator and discriminator use global self-attention across all layers to capture both short-range motifs and long-range dependencies. **(b)** Domain-adaptive pretraining on a curated RNAcentral corpus (333,627 sequences; ∼20M tokens), providing broad coverage of structural and regulatory non-coding RNAs for initializing RNAElectra representations. **(c)** Unified supervised fine-tuning of the same backbone across diverse downstream RNA tasks, spanning both classification and regression objectives. **(d)** BEACON task-level ranking summary across 13 benchmarks grouped into RNA structure, RNA function, and engineering categories. RNAElectra ranks first overall and places #1 on 9/13 tasks, indicating broad transfer across task families. **(e)** Radar plot summarizing task-wise BEACON performance, comparing RNAElectra with the strong RNA foundation model baseline RNA-FM (dashed line). The plot highlights consistent gains across heterogeneous endpoints (structure, function, engineering). For each task, the best-performing model among all compared methods is normalized to 100% and annotated as *Best in Class*, and RNA-FM is shown as a reference baseline.

To systematically evaluate transferability and robustness, we benchmarked RNAElectra on BEACON [11], a comprehensive suite of 13 RNA prediction tasks spanning structure inference, regulatory function, and engineering applications (Table 1). RNAElectra achieves the best overall performance on BEACON, attaining the top mean rank (1.96) and ranking first on 9 of 13 tasks (Fig. 1d). RNAElectra leads the full structure suite, including secondary structure prediction (SSP; F1 = 73.41%), structural score imputation (SSI; *R*^2^ = 48.02%), and tertiary-structure proxy tasks—contact map prediction (CMP; P@L = 74.14%) and distance map prediction (DMP; *R*^2^ = 56.90%). It also achieves best-in-benchmark performance on key functional and engineering endpoints, including alternative polyadenylation (APA; *R*^2^ = 89.81%), ncRNA classification (ACC = 97.29%), RNA modification prediction (Modif; AUC = 95.30%), programmable RNA switches (PRS; *R*^2^ = 58.87%), and vaccine degradation prediction (VDP; MCRMSE = 0.309) (Table 1). A head-to-head radar comparison with RNA-FM, a strong general-purpose RNA foundation model baseline, shows consistent gains across task categories (Fig. 1e). Across BEACON, RNAElectra also substantially exceeds naive supervised baselines trained from scratch (e.g., CNN/ResNet/LSTM), underscoring the value of RTD pretraining for learning transferable RNA regulatory representations (Table 1).

**Table 1.**
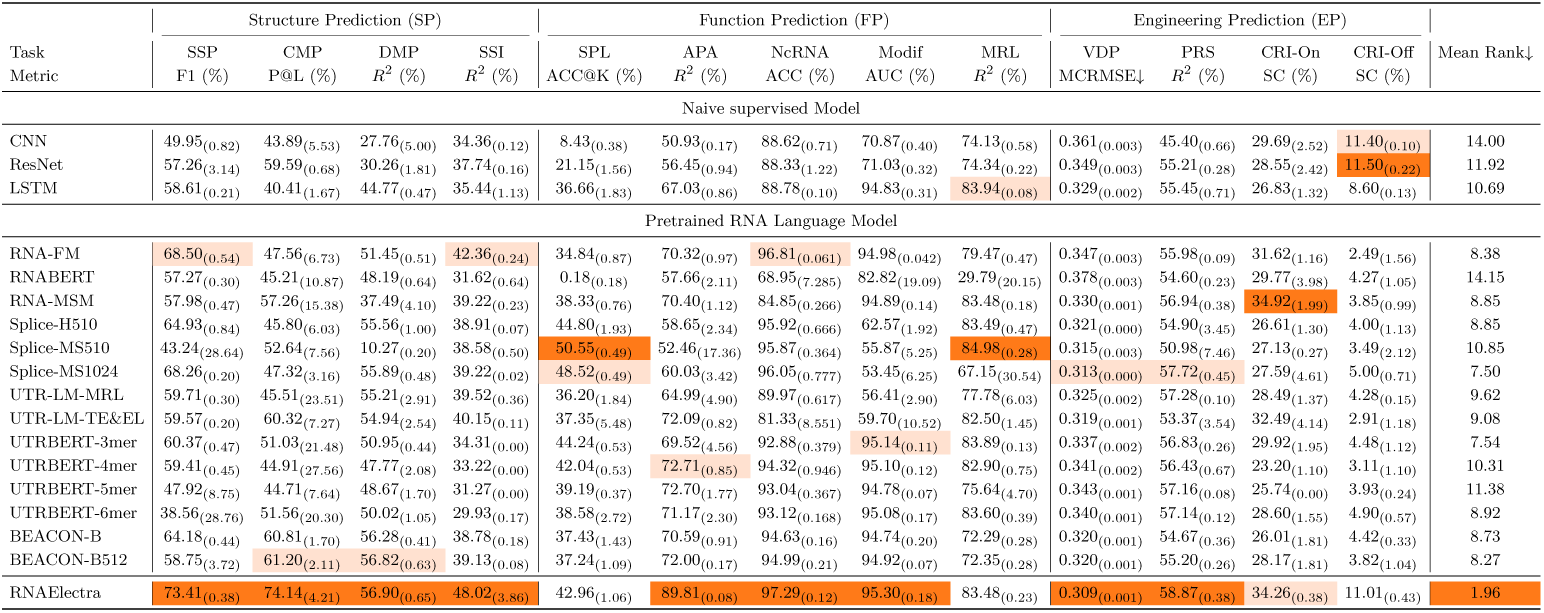
BEACON benchmark performance across 13 RNA prediction tasks spanning structure (SP), function (FP), and engineering (EP). The table summarizes how well models transfer from pretraining to diverse RNA endpoints under a unified evaluation protocol. Mean (s.d.) performance scores for baseline methods and previously reported pretrained RNA language models are reproduced from the BEACON study, while RNAElectra was evaluated on the same datasets, splits, and metrics. Best and second-best results per task are highlighted (higher is better unless noted); for vaccine degradation prediction (VDP), lower is better. The final column reports each model’s mean rank across all tasks, capturing overall breadth rather than performance on any single benchmark.

In the following sections, we analyze these results in depth, focusing on (i) embedding-space organization of ncRNA families, (ii) emergent structural sensitivity across secondary- and tertiary-structure proxies, and (iii) sequence determinants of RNA–protein binding, RNA modification, RNA–RNA targeting, translation, and stability.

### 2.2 RNAElectra embeddings cluster non-coding RNAs by functional class

Non-coding RNAs (ncRNAs) exhibit substantial functional diversity within the transcriptome, spanning catalytic ribozymes, translational repressors such as miRNAs, nucleolar guides including snoRNAs, and other specialized classes. Across these families, identity and function are rarely determined by a single signature; instead, they emerge from recurring combinations of short sequence motifs together with secondary-structure and contextual constraints that specify biogenesis pathways and molecular partners. We therefore asked whether RNAElectra learns representations that capture this family-level “grammar” and organize ncRNAs into coherent groups in embedding space without explicit supervision.

To test this, we visualized final-layer embeddings using t-SNE and quantified how well they support family discrimination across 21 annotated ncRNA families (as in RiNALMo [13]). RNAElectra produces compact, well-separated family-level structure in embedding space (Fig. 2b) and achieves the highest *embedding-based* Macro F1 (0.997), slightly exceeding RNA-FM and RiNALMo (both 0.995), whereas RNAErnie and SpliceBERT show weaker separation.

**Fig. 2.**
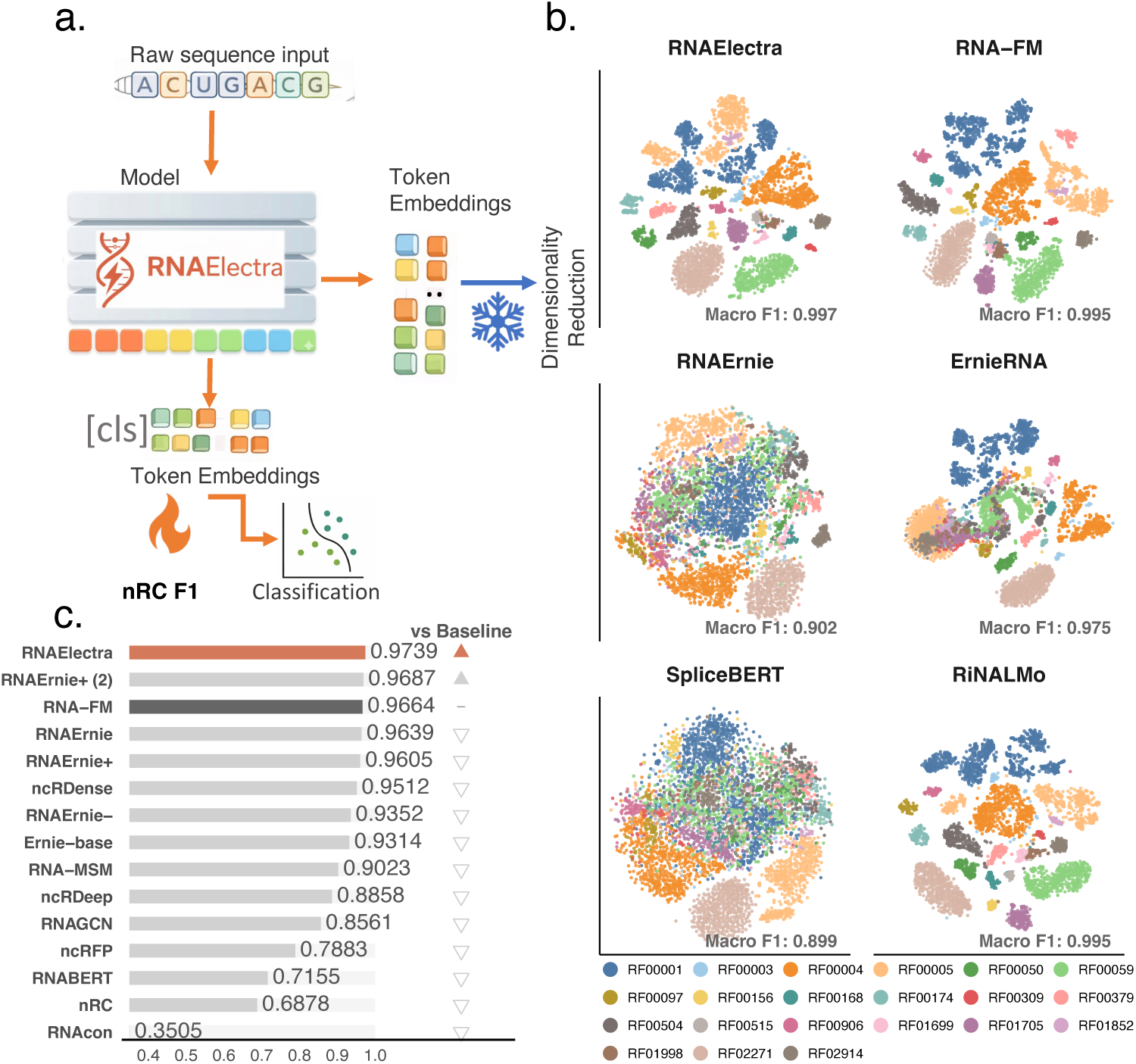
RNAElectra embeddings separate ncRNA families and support ncRNA functional classification. **(a)** Evaluation workflow: RNAElectra encodes an RNA sequence into contextual token embeddings; the `[CLS]` embedding is used as a sequence representation for down-stream classification, and embeddings are projected to low dimensions for visualization. **(b)** t-SNE projections of final-layer embeddings for 21 annotated ncRNA families, comparing RNAElectra with representative baseline RNA language models. RNAElectra forms compact, well-separated family-level clusters and achieves the highest clustering score (Macro F1 = 0.997), marginally above RNA-FM and RiNALMo (both 0.995), whereas RNAErnie and SpliceBERT exhibit weaker separation. **(c)** Supervised ncRNA family classification on the nRC benchmark [28]. Bars report F1-scores for RNAElectra and comparison methods; RNAElectra ranks first overall (F1 = 0.9739).

We next asked whether these representations translate into improved supervised performance. On ncRNA family classification using the nRC benchmark [28], RNAElectra ranks first overall and attains the best F1-score (0.9739), outperforming RNA-FM (0.9664), the strongest RNAErnie variants (up to 0.9687), and graph-based and classical baselines (Fig. 2c). Together, these results indicate that RNAElectra learns sequence representations that capture discriminative ncRNA family signatures and support state-of-the-art performance for ncRNA annotation.

### 2.3 RNAElectra learns emergent RNA structural constraints

RNA exerts its function through folding, where secondary structure—local helices and loops—provides a scaffold for tertiary interactions that enable binding, catalysis, and regulatory control. We asked whether a sequence-only foundation model can recover structural information as a consequence of pretraining. Because RNAElectra is trained to detect context-dependent nucleotide replacements across the entire sequence, it is encouraged to learn which substitutions remain compatible with surrounding context—constraints shaped by base-pairing patterns, loop composition, and longer-range dependencies.

Across the BEACON structure suite (SSP, SSI, CMP, DMP), RNAElectra shows the strongest overall profile among general-purpose RNA language models (Fig. 3a). On secondary structure prediction (SSP), RNAElectra achieves the top performance (F1 = 73.41%), improving over RNA-FM (68.50%) by 4.91 points (Fig. 3b). This advantage extends beyond BEACON: RNAElectra attains F1 = 0.933 on ArchiveII600 (precision 0.960, recall 0.918) and F1 = 0.715 on TS0, exceeding RNA-FM and other sequence-only baselines on both benchmarks (Fig. 3c; Table S4). Improvements extend to tertiary-structure proxy tasks. RNAElectra achieves the best performance on structural score imputation (SSI; *R*^2^ = 48.02%), outperforming RNA-FM (*R*^2^ = 42.36%) and UTR-LM-TE&EL (*R*^2^ = 40.15%) (Fig. 3d). It also leads on contact-map prediction (CMP; P@L = 74.14%) and distance-map prediction (DMP; *R*^2^ = 56.90%), surpassing RNA-FM (CMP P@L = 47.56%, DMP *R*^2^ = 51.45%) and other general-purpose models (Fig. 3e i, ii).

**Fig. 3.**
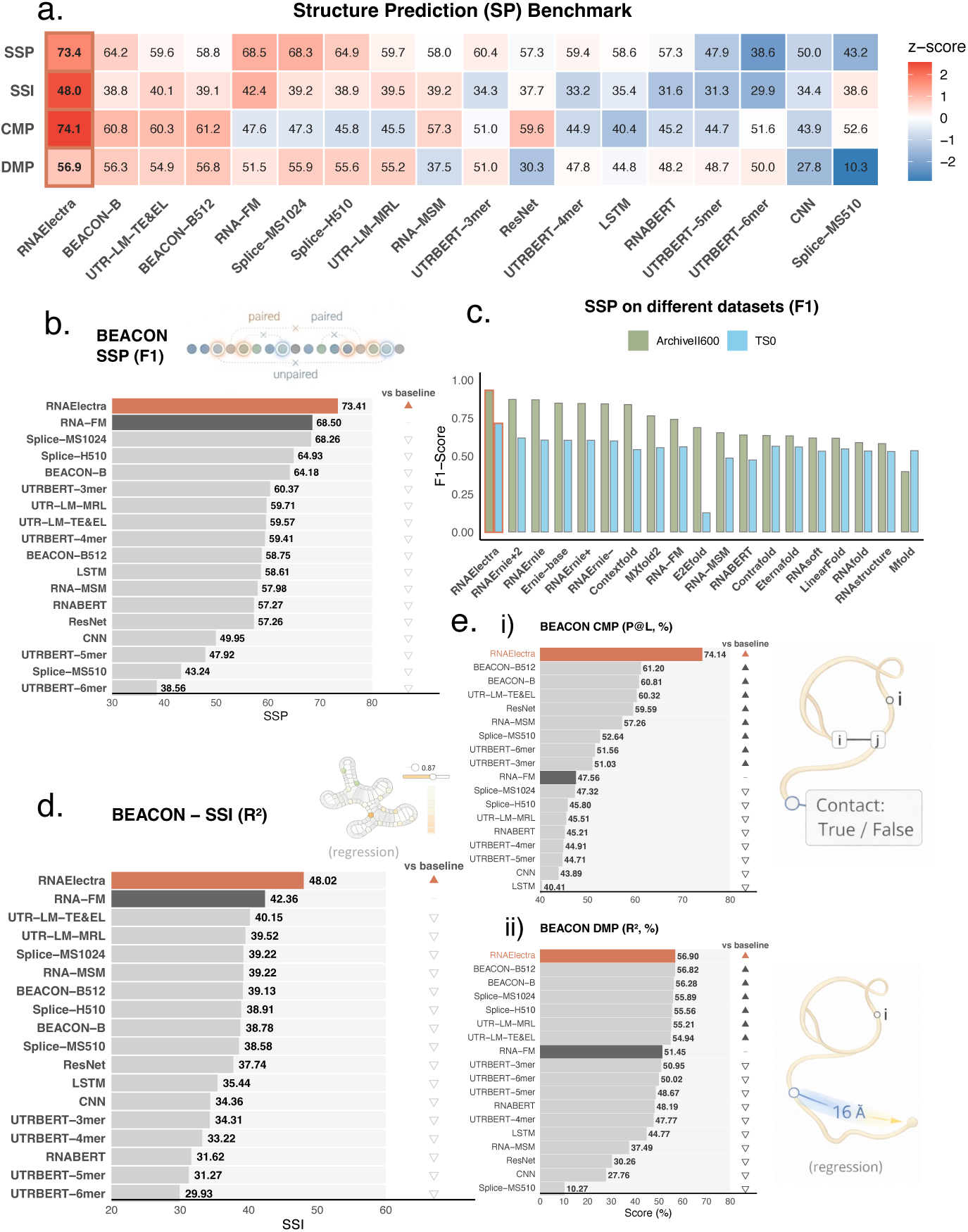
RNAElectra achieves state-of-the-art performance on RNA structure–related tasks. **(a)** BEACON structure prediction (SP) overview across secondary structure prediction (SSP), structural score imputation (SSI), contact map prediction (CMP), and distance map prediction (DMP). Cells report z-score–normalized performance with raw scores overlaid; RNAElectra ranks first on all four tasks. **(b)** SSP on BEACON (F1). RNAElectra attains the best score (F1 = 73.41%) compared with the strong general-purpose RNA foundation baseline RNA-FM (68.50%) and other pretrained and supervised baselines; triangles summarize changes relative to RNA-FM. **(c)** SSP on independent datasets (ArchiveII600 and TS0). RNAElectra attains F1 = 0.933 on ArchiveII600 and F1 = 0.715 on TS0, exceeding RNA-FM and remaining competitive with structure-specialized baselines across both datasets. **(d)** Structural Score Imputation (SSI; *R*^2^) on BEACON, with RNAElectra achieving the highest regression performance. **(e)** Tertiary-structure proxy tasks on BEACON: (i) CMP (P@L) and (ii) DMP (*R*^2^). RNAElectra ranks first on both CMP (P@L = 74.14%) and DMP (*R*^2^ = 56.90%) relative to RNA-FM and other compared models.

Overall, RNAElectra captures structure-relevant constraints directly from sequence, recovering both local pairing preferences and longer-range compatibility without using structural annotations, alignments, or probing measurements as input features. This sensitivity is particularly useful in sequence-only settings, where folding-relevant signals must be inferred from primary sequence alone.

### 2.4 RNAElectra captures the sequence binding rules of RNA-Protein, modification, and RNA-RNA interactions

Once folded, RNAs engage in dynamic interactions with proteins, chemical modifications, and other RNAs, forming core layers of post-transcriptional regulation. These interactions are often specified by compact sequence codes—short binding motifs for RNA-binding proteins (RBPs), local consensus contexts for modifications, and base-pairing rules for small-RNA targeting—embedded within broader sequence context. We asked whether RNAElectra, trained purely on sequence, learns representations that generalize across these heterogeneous interaction regimes.

Across a large-scale RBP binding benchmark comprising 313 CLIP-seq experiments across 203 RBPs [29], RNAElectra achieves the strongest overall performance among sequence-only models, with mean AUROC of 0.9068 under Neg-1 and 0.8570 under the more stringent Neg-2 setting (Fig. 4a). These two regimes probe distinct failure modes: Neg-1 contrasts true binding sites against non-RBP background sequences, whereas Neg-2 requires distinguishing true sites from binding sites of other RBPs, a harder setting where negatives can share similar composition and motif content. Notably, high accuracy is paired with stability across regimes. When plotting Neg-2 AUROC against robustness (1−Δ, where Δ is the AUROC drop from Neg-1 to Neg-2), RNAElectra lies in the high-performance, high-robustness region and is the top sequence-only method on both axes (Fig. 4a; Δ = −0.0498). While models that incorporate additional region features (e.g., transcript segment annotations such as 5^′^UTR/CDS/3^′^UTR) and/or explicit structural inputs can attain higher absolute AUROC, RNAElectra consistently leads the sequence-only group, indicating that a substantial fraction of RBP-binding signal is recoverable from sequence context alone.

**Fig. 4.**
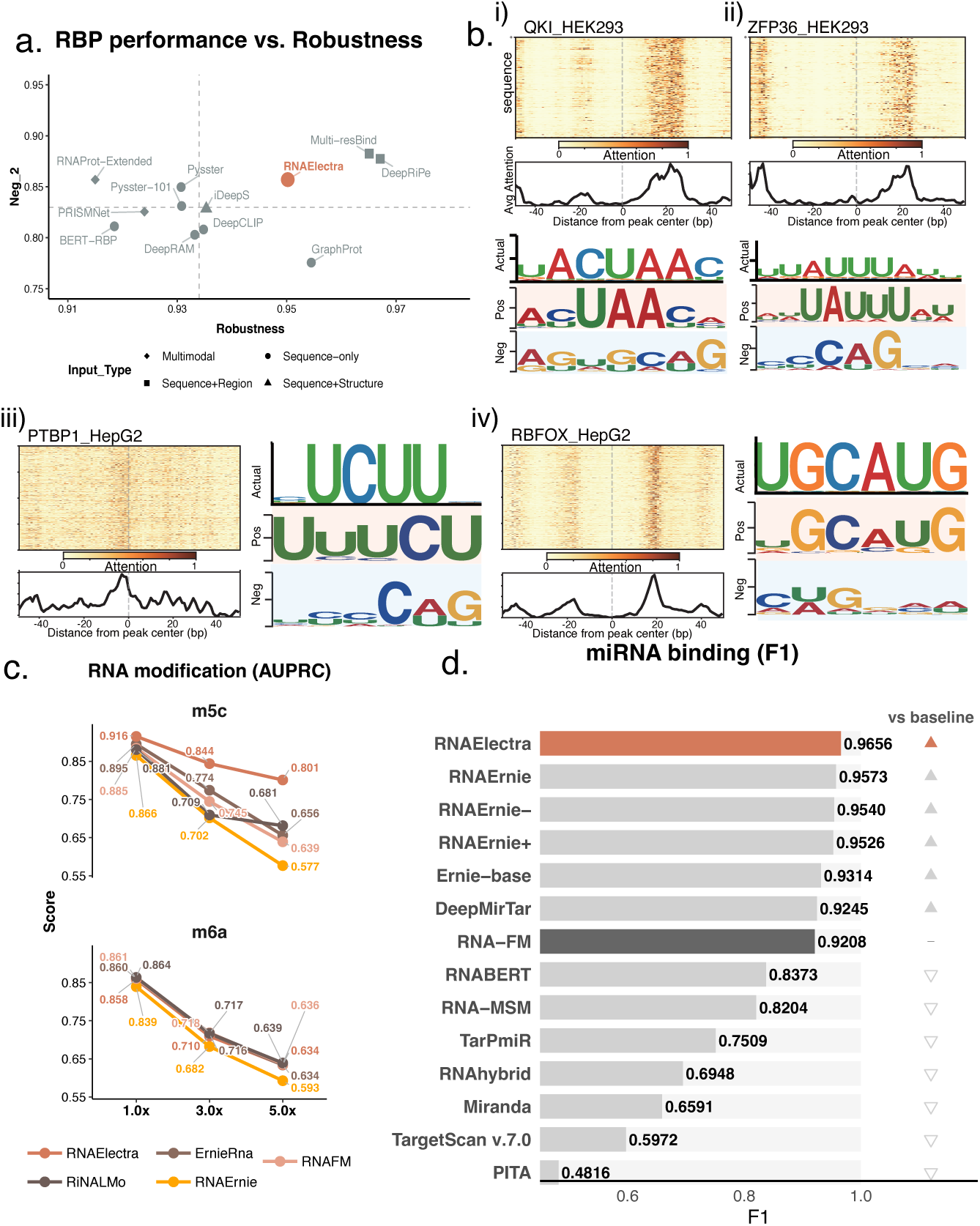
RNAElectra captures multiple RNA interaction codes across protein binding, RNA modification, and RNA–RNA targeting. **(a)** RBP binding performance versus robustness across models. The y-axis reports Neg-2 AUROC and the x-axis reports robustness (1−Δ), where Δ is the AUROC drop from Neg-1 to Neg-2; RNAElectra achieves a strong balance of accuracy and robustness among sequence-only and feature-augmented approaches. **(b)** Motif-centric interpretability for representative RBPs. For QKI (HEK293), ZFP36 (HEK293), PTBP1 (HepG2), and RBFOX (HepG2), peak-centered attention heatmaps and averaged attention profiles localize model signal near binding centers. Sequence logos recovered from high-scoring positive predictions recapitulate known RBP motifs, whereas logos from predicted negative sets do not recover the canonical motif patterns. **(c)** RNA modification prediction for m^5^C and m^6^A under two class ratios (1:1 and 1:5 positives:negatives). RNAElectra achieves strong performance on m^5^C and competitive performance on m^6^A relative to representative RNA foundation model baselines. **(d)** miRNA–target interaction prediction on the DeepMirTar benchmark, evaluated by F1 score; RNAElectra achieves the top performance (F1 = 0.9656) among compared methods.

To connect performance to sequence determinants, we performed motif analysis around predicted binding sites for representative RBPs (QKI, ZFP36, PTBP1, and RBFOX; Fig. 4b). Peak-centered attention heatmaps and average attention profiles concentrate around binding centers, and sequence logos from motif enrichment on high-scoring positive predictions recapitulate known RBP motifs, whereas logos derived from predicted negative sets do not recover the canonical motif patterns (Fig. 4b).

We next assessed whether the same sequence representations support prediction of covalent RNA marks. Using m^6^A-Atlas v2.0 [30] and m^5^C-Atlas [31] as reference resources, we evaluated modification-site prediction using AUPRC under increasing class imbalance (1×, 3×, and 5× negatives; Fig. 4c). RNAElectra achieves the best performance across all settings for m^5^C (AUPRC = 0.916/0.844/0.801 at 1×/3×/5×), and remains competitive for m^6^A across ratios (AUPRC = 0.861/0.718/0.636), comparable to the strongest baselines as imbalance increases.

Finally, we evaluated miRNA target-site prediction on DeepMirTar [32], which aggregates *>*14,000 experimentally supported sites from CLASH and curated databases. RNAElectra achieves the highest F1-score (0.9656), outperforming widely used RNA foundation model baselines (e.g., RNA-FM and RNAErnie) as well as a strong task-specific neural baseline (DeepMirTar) (Fig. 4d). In this setting, performance depends on capturing constrained miRNA–mRNA base-pairing rules and local target-site context. Notably, RNAElectra and other modern RNA foundation models substantially outperform widely used classical tools (e.g., TargetScan and miRanda) on DeepMirTar (Fig. 4d), consistent with foundation models capturing informative sequence determinants of miRNA targeting.

Together, these results show that beyond fold-relevant constraints, RNAElectra captures multiple interaction codes—including RNA–protein binding motifs, sequence contexts associated with RNA modifications, and RNA–RNA targeting signals— yielding strong and robust sequence-only performance across complementary layers of post-transcriptional regulation.

### 2.5 RNAElectra captures quantitative determinants of translation efficiency and mRNA stability

Once an RNA is transcribed, its sequence must satisfy multiple quantitative constraints: recruit and position ribosomes to set protein output, and encode decay-linked signals that determine how long the transcript persists. Unlike discrete family labels or binding-site calls, these phenotypes are continuous readouts that integrate many weak cues distributed across the molecule. We therefore asked whether a sequence-only foundation model—trained without experimental measurements—can recover these graded regulatory programs directly from sequence.

We evaluated RNAElectra on three complementary regression settings that probe distinct layers of post-transcriptional control: mRNA stability/half-life from large reporter compendia, translation efficiency (TE) from matched Ribo-seq/RNA-seq, and mean ribosome loading (MRL) from massively parallel 5^′^UTR reporter assays (Fig. 5a).

**Fig. 5.**
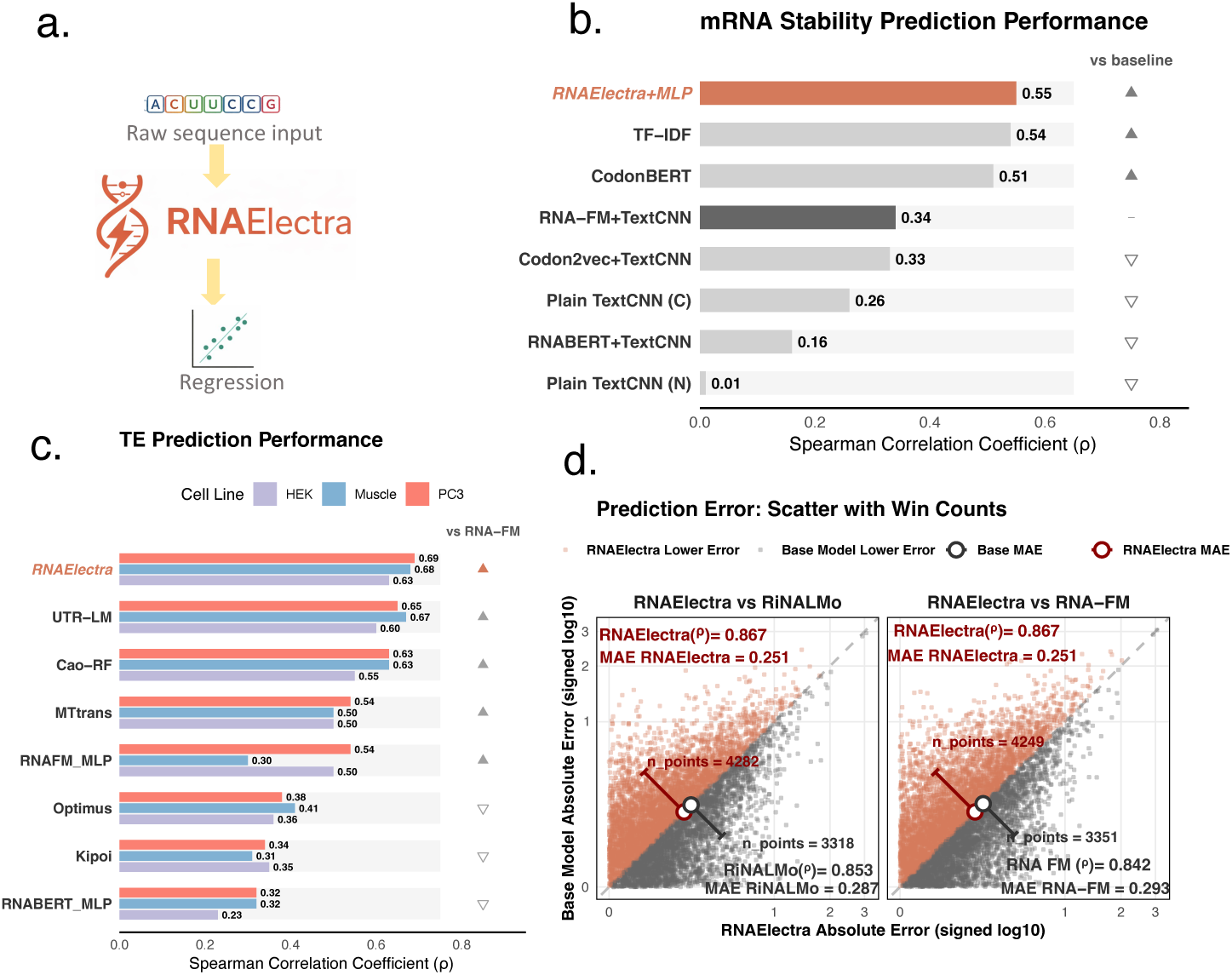
RNAElectra captures quantitative determinants of mRNA stability and translation. **(a)** Regression evaluation setting: RNAElectra encodes the input RNA sequence and a regression head predicts a continuous regulatory readout. **(b)** mRNA stability prediction measured by Spearman correlation (*ρ*). RNAElectra+MLP achieves the top performance (*ρ* = 0.55), exceeding all compared codon-based and nucleotide-based models. **(c)** Translation efficiency (TE) prediction measured by Spearman correlation (*ρ*) across HEK293T, Muscle, and PC3. RNAElectra achieves the highest correlations across cell lines (0.63–0.69), outperforming RNA-FM and other baselines. **(d)** Mean ribosome loading (MRL) prediction error comparison against RiNALMo and RNA-FM. Scatter plots show per-transcript absolute prediction errors of RNAElectra (x-axis) versus the baseline model (y-axis; RiNALMo or RNA-FM), displayed under a signed log_10_ transformation to emphasize both small and large deviations. Points are colored by which model achieves lower error (RNAElectra in red, baseline in dark gray, ties in light gray). Open circles denote mean absolute error (MAE) for each model, and annotated counts report the number of transcripts where each model attains lower error. Spearman correlation (*ρ*) between predictions and measurements is shown for each model; RNAElectra achieves higher *ρ* and lower MAE, and attains lower error for more transcripts than either baseline.

We first tested whether RNAElectra captures sequence programs associated with mRNA lifetime, a phenotype shaped by distributed cis-elements and context-dependent decay pathways. On a large mRNA stability benchmark (41,123 transcripts spanning broad sequence lengths), RNAElectra achieves the highest correlation with measured stability (Spearman *ρ* = 0.55), outperforming both codon-based and nucleotide-based baselines (Fig. 5b). This performance is obtained without codon tokenization or engineered features, suggesting that RNAElectra learns informative sequence determinants associated with mRNA decay.

We next evaluated whether RNAElectra captures sequence determinants of translation efficiency measured from matched Ribo-seq/RNA-seq. Across endogenous TE datasets (HEK293T, Muscle, and PC3), RNAElectra achieves strong correlations with measured TE (Spearman *ρ* = 0.63–0.69) and performs among the best models across cell lines (Fig. 5c). Performance is consistent across cell lines, supporting transfer of sequence determinants of TE using sequence input alone.

Finally, we evaluated RNAElectra on massively parallel reporter assays for mean ribosome loading (MRL), where standardized constructs isolate the contribution of 5^′^UTR sequence to translation. RNAElectra achieves the highest agreement with measured MRL among the compared foundation models (Spearman *ρ* = 0.867, versus RiNALMo *ρ* = 0.853 and RNA-FM *ρ* = 0.842). To further characterize error patterns, we compared per-transcript absolute errors in pairwise scatter plots (Fig. 5d), where RNAElectra shows lower error for a larger fraction of transcripts and a lower mean absolute error (MAE = 0.251 vs. 0.287 and 0.293 for RiNALMo and RNA-FM, respectively).

Together, these regression results complement the classification and interaction benchmarks by showing that RNAElectra captures transferable, quantitative sequence determinants spanning translation-related readouts and transcript stability under a unified, sequence-only modeling framework.

## 3 Discussion

RNAElectra demonstrates that ELECTRA-style replaced-token detection (RTD) is an effective and practical pretraining objective for RNA foundation modeling at single-nucleotide resolution. Across diverse prediction settings—including structure-related tasks, RNA–protein and RNA–RNA interactions, RNA modification landscapes, and quantitative regulatory readouts—RNAElectra exhibits robust cross-task transfer under a unified fine-tuning protocol. These results support a broader point: because many RNA functions are encoded by weak, distributed sequence cues, objectives and tokenization schemes that provide dense, position-wise learning signals can improve downstream generalization and facilitate model interpretation. The distribution of gains across benchmarks is consistent with RTD preferentially benefiting *site-centric* tasks whose decision boundaries are defined by sharp, nucleotide-level rules (e.g., motif cores and their immediate context). In contrast, *aggregate* readouts over longer regions—including translation-related phenotypes or stability/half-life—can depend more strongly on broader sequence composition, long-range context, and dataset-specific factors (e.g., sequence length distributions, negative sampling, and measurement protocols), which can reduce the marginal advantage of any single objective. This perspective helps interpret where dense, discriminator-based supervision is most impactful and where additional inductive biases may be beneficial.

### RTD aligns pretraining with RNA sequence-to-function inference

A central distinction between RNAElectra and most existing RNA foundation models is the pretraining objective. Masked language modeling (MLM) concentrates supervision on a small subset of positions and optimizes on masked inputs that are absent at down-stream inference, whereas RTD trains a discriminator to judge token authenticity at every position in the context of a fully observed (but realistically corrupted) sequence. By defining the loss over all positions, RTD provides dense, position-wise learning signals and encourages sensitivity to subtle, context-dependent deviations throughout the sequence, plausibly benefiting settings where single-nucleotide changes can produce large functional effects. This is particularly relevant for RNA, where regulatory mechanisms such as RBP binding and modification contexts are local yet strongly modulated by surrounding sequence. A distinctive feature of RTD is that the discriminator must distinguish *authentic* nucleotides from *plausible replacements* proposed by the generator, explicitly training the model to separate authentic sequence contexts from close “near-match” variants rather than only reconstructing masked tokens. For example, functional sites are often surrounded by near-matches—a canonical miRNA seed match with a single mismatch or bulge, or an RBP motif instance with one base substituted—that are difficult to distinguish without modeling local context and broader compatibility. Consistent with this, RNAElectra shows strong performance and robustness in settings with harder negatives (e.g., Neg-2 RBP binding, where negatives are binding sites of other RBPs) and recovers canonical motif cores from high-confidence predictions. Moreover, RTD provides a model-native, position-wise signal: the discriminator outputs a per-position replacement probability that can be interpreted as an *authenticity* (plausibility) score for each nucleotide given its context. This type of per-nucleotide plausibility score is less direct in MLMs, which typically require pseudo-likelihood evaluation via repeated masking.

### Nucleotide-resolution representations support RNA regulatory grammar and long-range context

RNA regulatory mechanisms are often governed by base-level rules—motif cores for RBPs, modification-context preferences, and explicit base pairing in small-RNA targeting—so nucleotide-resolution representations provide a natural interface for interpretation and sequence editing. Models that operate on *k*-mers can be effective for coarse discrimination, but may blur single-nucleotide effects that matter for regulatory interpretation and rational editing. RNAElectra models sequences at nucleotide resolution, enabling fine-grained attribution and reducing ambiguity when linking predictions to candidate determinants. At the same time, RNA function is not purely local: longer-range dependencies arise through folding, cooperative binding, and broader transcript context, making efficient global context integration important for tasks that combine motif-scale signals with distal context. Finally, relative to genome-scale DNA contexts, many RNA benchmarks operate on shorter sequences, making nucleotide-level Transformers computationally tractable while still benefiting from global context modeling.

### Cross-task transfer is evaluated under a unified, sequence-only fine-tuning protocol

A primary value of RNA foundation models is the ability to reuse a single pretrained backbone encoder across heterogeneous tasks without bespoke engineering. RNAElectra is evaluated using a unified fine-tuning protocol that avoids task-specific architectural modifications or auxiliary inputs and keeps downstream heads and training procedures consistent when comparing different pretrained encoders. This provides a controlled setting for representation-centric comparisons and reduces the likelihood that improvements are driven primarily by task-specific head design or feature pipelines, while emphasizing portability across datasets and experimental settings where consistent side information may be unavailable.

### Interpretability analyses connect predictions to sequence determinants

Beyond predictive benchmarking, RNA applications often benefit from interpretability to support hypothesis generation and sequence-level reasoning. We therefore complement performance evaluations with targeted analyses—including embedding-space organization of ncRNA families and motif discovery from high-confidence RBP binding predictions—to connect model outputs to plausible sequence determinants. Importantly, single-nucleotide tokenization improves interpretability by enabling base-resolved attribution, making it easier to localize which positions and local contexts contribute most to a prediction. While such analyses do not establish causality *in vivo*, they provide practical evidence that the model captures biologically meaningful signals and help identify where predictions are most sensitive to sequence context.

In summary, RNAElectra advances RNA foundation modeling by pairing RTD pre-training with nucleotide-resolution representations and efficient context integration, yielding a practical backbone that transfers across diverse RNA prediction tasks under a unified evaluation protocol. These findings motivate broader exploration of dense, discriminator-based objectives for biological sequence modeling and provide a foundation for future work integrating additional priors and expanding robustness evaluation across datasets and experimental settings.

## 4 Conclusions

We presented RNAElectra, an RNA foundation model that advances RNA sequence modeling by combining single-nucleotide resolution with an ELECTRA-style replaced-token detection (RTD) pretraining objective and an efficient global attention design. Pretrained on diverse non-coding RNAs, RNAElectra provides a practical and reusable backbone encoder for RNA prediction that operates in a unified, sequence-only setting without task-specific architectures or auxiliary inputs. Across broad benchmarking spanning RNA structure and function, RNA–protein and RNA–RNA interactions, RNA modification landscapes, and quantitative regulatory readouts (including translation efficiency and mRNA stability), RNAElectra outperforms widely used RNA foundation model baselines on the majority of evaluated tasks while remaining competitive elsewhere. These results support a central conclusion: dense, position-wise discriminator training is an effective alternative to masked language modeling for RNA foundation modeling, better aligning pretraining with downstream inference on fully observed sequences and improving cross-task transfer for diverse RNA regulatory problems. Beyond predictive performance, RNAElectra enables sequence-level inspection through analyses that relate learned representations to underlying sequence determinants, helping connect model outputs to biological hypotheses and sequence design. Collectively, RNAElectra establishes RTD pretraining as a compelling foundation-model objective for RNA and provides a scalable framework for future extensions that incorporate additional data diversity and regulatory contexts to further broaden generalization and utility.

## 5 Materials and Methods

### 5.1 Training strategies and pretraining objective

#### Overview

RNAElectra follows a two-stage pipeline. We first pretrain *from scratch* on a curated RNAcentral corpus using an ELECTRA-style generator–discriminator objective to learn general-purpose RNA sequence representations. We then fine-tune the pretrained discriminator under a unified protocol across diverse downstream RNA tasks to assess transfer to structure, interaction, and regulatory prediction settings.

#### Replaced-token detection (RTD) pretraining

RNAElectra is pretrained using an ELECTRA-style replaced-token detection (RTD) objective with a generator– discriminator framework. Given an input sequence *X* = {*x*_1_*,…, x_N_* }, we sample a set of masked positions *M* (mask rate *r*) and form *X*_mask_ by replacing *x_i_* with `[MASK]` for all *i* ∈ *M*. The generator *G* is trained with masked language modeling (MLM) to recover the original nucleotides at masked positions:

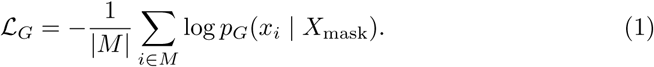

We then construct a corrupted sequence *X*^′^ by replacing each masked position *i* ∈ *M* with a nucleotide sampled from *p_G_*(· | *X*_mask_), while leaving all unmasked positions unchanged. To avoid ambiguity in RTD labels, we *disallow same-token replacements*: if the sampled nucleotide equals the original nucleotide *x_i_*, we resample until a different nucleotide is drawn. The discriminator *D* takes *X*^′^ as input and outputs, for each position *i*, a probability *ŷ_i_* = *p_D_*(*y_i_*= 1 | *X*^′^) that the token at position *i* is *original* (not replaced). With the above corruption procedure, RTD labels can be defined equivalently by token identity: *y_i_* = 1 if *x*^′^_*i*_ = *x_i_* and *y_i_*= 0 otherwise. The discriminator is trained with binary cross-entropy over all positions:

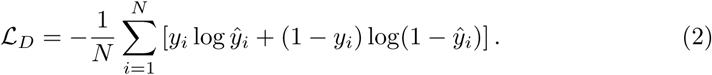

The total pretraining loss is

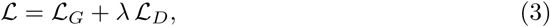

where *λ* controls the relative weight of RTD and generator MLM. We use dynamic masking during pretraining such that masked positions *M* are resampled each step.

#### Masking and corruption policy

We use dynamic masking during pretraining such that masked positions are resampled each step. Unless otherwise specified, we mask a fraction *r* = [0.15] of positions. For the generator replacements, we use argmax from *p_G_*(· | *X*_mask_), and allow replacing a position with its original nucleotide. We set *λ* = [50]. Full optimizer, schedule, and compute details are provided in Appendix S1.

### 5.2 Model architecture

RNAElectra implements an ELECTRA generator–discriminator framework. The *generator* is a lightweight Transformer with 12 layers and hidden size 256, trained with MLM to propose nucleotide replacements for masked positions. The *discriminator* is the backbone used for downstream tasks; it contains 22 Transformer layers with hidden size 512 and predicts, at each position, whether the token is original or replaced.

#### Efficient attention for long RNAs

To improve scalability to long RNA sequences, we implement full-sequence global self-attention using FlashAttention-2, which provides a memory- and compute-efficient realization of attention for long inputs while preserving modeling capacity for long-range interactions.

#### Implementation details

Unless otherwise specified, we use standard Transformer components with 16 attention heads and a feed-forward width of 2048. We do not share token embeddings between generator and discriminator. Additional architectural and training hyperparameters are reported in Supplementary S1.

### 5.3 Pre-training data and tokenization

#### RNAcentral pretraining corpus

The pretraining corpus was derived from the RNAcentral Active collection (integrating Ensembl, Rfam, RefSeq). We removed sequences containing ambiguous nucleotide N and truncated sequences to a maximum length of 1024 nucleotides (discarding sequences shorter than 10 nucleotides). The resulting dataset was randomly split into training (90%) and validation (10%).(details in Supplementary S2).

#### Single-nucleotide tokenization

Sequences are tokenized at single-nucleotide resolution, treating each base as one token. We use an {A,C,G,U} nucleotide alphabet and map uracil (U) to thymine (T) to obtain an {A,C,G,T} representation for tokenization, yielding a compact nucleotide vocabulary while preserving nucleotide-level information. We add standard special tokens (e.g., `[CLS]` for sequence-level prediction and `[MASK]` for generator MLM) following common Transformer practice. This representation supports nucleotide-resolution predictions required by RNA regulatory tasks, including modification-site and binding-site identification.

### 5.4 Downstream tasks fine-tuning details

#### Training setup

To enable fair and reproducible comparisons, we adopt the BEACON benchmark fine-tuning protocol and keep the supervised training configuration fixed across all evaluated pretrained backbones. Unless otherwise specified, we *fully fine-tune* all model parameters (no frozen layers) and use the same optimizer, schedule, and data splits as BEACON. For each model–task pair, we sweep the learning rate in the range [1 × 10^−5^, 5 × 10^−3^] and select the best setting based on validation performance. All experiments are repeated with three random seeds, and we report the mean performance together with the sample standard deviation. Additional implementation details (including random seeds, batch sizes, and learning rates) are provided in Supplementary S3.

#### Task pipeline

Following BEACON, we implement three prediction pipelines to accommodate different output granularities: *sequence-level*, *nucleotide-level*, and *nucleotide–nucleotide relation* prediction. RNA structural readouts with structured outputs—*Secondary Structure*, *Contact Map*, and *Distance Map*—are formulated as nucleotide–nucleotide relation prediction problems (within-sequence pairing or distance/contact prediction, depending on the task).

#### Sequence-level prediction

For sequence-level tasks, we obtain a single embedding per input sequence. For Transformer-based language models, we use the contextual representation of the `[CLS]` token. The sequence embedding is passed to a linear classification head to produce the final prediction.

#### Nucleotide-level prediction

For nucleotide-level tasks, we output one prediction per nucleotide position by applying an MLP head to each contextualized nucleotide embedding. For the SSI task specifically, each nucleotide embedding is concatenated with a projected structural score feature before being passed to the prediction head. For tokenizations that span multiple nucleotides, we compute a nucleotide embedding by averaging the embeddings of all tokens that cover that position, ensuring a consistent nucleotide-resolution output interface across models.

#### Nucleotide–nucleotide relation prediction

For within-sequence structure tasks (SSP, CMP, DMP), we formulate prediction as a nucleotide–nucleotide pairing problem within a single RNA sequence of length *l*. We encode the sequence to obtain contextualized nucleotide embeddings **H** ∈ ℝ*^l^*^×*d*^, then construct pairwise features for every nucleotide pair (*i, j*) with *i, j* ∈ [1*, l*] via outer concatenation of **H** with itself, assembling an *l* × *l* interaction map. This map is processed by a lightweight 2D residual network (ResNet) head comprising 16 residual blocks with 64 channels and 3 × 3 convolutions to model local 2D patterns, producing final pairing scores or distance estimates.

### 5.5 BEACON Benchmark Tasks

We evaluate RNAElectra on BEACON, a comprehensive RNA benchmark comprising 13 tasks spanning *structure*, *function*, and *engineering* applications, with standardized data splits, evaluation metrics, and a unified fine-tuning protocol [11]. BEACON includes both sequence-level and nucleotide-level settings, and additionally formulates structured structure-related outputs (e.g., pairing/contact/distance maps) as nucleotide–nucleotide relation prediction tasks [11]. Unless otherwise specified, we follow the task definitions, dataset construction, and metrics provided in the BEACON benchmark.

#### Task groups

Briefly, the structural analysis tasks include secondary structure prediction (SSP), contact map prediction (CMP), distance map prediction (DMP), and structural score imputation (SSI). Functional tasks include splice site prediction (SPL), alternative polyadenylation isoform prediction (APA), non-coding RNA function classification (ncRNA), RNA modification prediction (Modif), and mean ribosome loading prediction (MRL). Engineering-oriented tasks include vaccine degradation prediction (VDP), programmable RNA switches (PRS), and CRISPR on-target/off-target prediction (CRI-On/CRI-Off) [11].

#### Outputs and metrics

Across these tasks, BEACON evaluates classification and regression settings using task-appropriate metrics (e.g., F1 for SSP, Top-*L* precision for CMP, *R*^2^ for regression tasks, AUC for imbalanced multi-label prediction, MCRMSE for VDP, and Spearman correlation for CRISPR tasks) [11]. A complete summary of dataset sizes, sequence lengths, sources, and metrics is provided by BEACON (Table 1 and Sec. 3) [11].

### 5.6 Datasets for extended downstream tasks

#### RNA secondary structure data

To assess generalization in RNA secondary structure prediction beyond BEACON, we follow the evaluation protocol used by RNAErnie [18]. Specifically, we (i) train on RNAStralign [33] and evaluate on ArchiveII600 (a length-restricted subset of ArchiveII [34]), and (ii) train/validate/test on the bpRNA-1m splits TR0/VL0/TS0 introduced in SPOT-RNA [35]. ArchiveII600 is a widely used benchmark comprising curated reference secondary structures from multiple RNA families, with all sequences shorter than 600 nucleotides. TS0 is an independent, non-redundant test split constructed from bpRNA with CD-HIT-EST filtering (80% identity) and a TR0/VL0/TS0 partition.

*RNAStralign [33]:* RNAStralign contains 37,149 RNA secondary structures from eight RNA families with lengths of approximately 100–3,000 nt. After applying the 600-nt cutoff for training, we retain 20,923 structures across the same eight families.

*ArchiveII / ArchiveII600 [34]:* ArchiveII includes 3,975 RNA structures from ten RNA families with lengths of approximately 100–2,000 nt. ArchiveII600 denotes the subset with sequence length *<* 600 nt, which we use for evaluation.

*bpRNA-1m TR0/VL0/TS0 [35]:* bpRNA-1m contains 13,419 non-redundant RNA secondary structures spanning 2,588 families, obtained by removing redundancy with CD-HIT-EST (80% identity) and splitting the resulting set into TR0 (10,814), VL0 (1,300), and TS0 (1,305).

We report base-pair prediction quality using precision, recall, and F1 computed between predicted and reference base pairs (see Supplementary Table S4).

#### Protein–RNA interaction data (RBP benchmark)

The benchmark integrates three CLIP collections: ENCODE eCLIP (223 experiments across 150 RBPs in HepG2/K562), iONMF (31 experiments across 19 RBPs spanning multiple CLIP protocols), and the Mukherjee PAR-CLIP compendium (59 experiments in HEK293), totaling 313 experiments across 203 RBPs [29]. Inputs are defined around single-nucleotide crosslink sites (e.g., peak centers / 5^′^ peak ends depending on the source dataset) by extracting fixed-length sequence windows (50nt) centered at each site. [29] For each experiment, the benchmark provides two negative-control schemes: *neg-1*, sampled uniformly from positions on transcripts that overlap at least one binding site of the target RBP, and *neg-2*, sampled from binding sites of other RBPs in the same dataset (restricted to non-overlapping sites to avoid co-binding leakage), with negatives generated at a 1:1 ratio to positives and models evaluated separately under each scheme. [29] To align with the RBP benchmark, the mean AUROC is reported for two negative-control schemes. (see Supplementary Table S5)

#### Epitranscriptomic modification data

##### Data sources

For modification-related analyses, we used curated atlas resources including m^6^A-Atlas v2.0 [30] and m^5^C-Atlas [31], which aggregate high-confidence sites from multiple experimental protocols and biological contexts. For m^6^A, the atlas integrates sites supported by 12 profiling techniques (including miCLIP, m6A-SAC-seq, MAZTER-seq, m6A-REF-seq, m6A-CLIP-seq, DART-seq, and related assays), spanning 22 cell lines/tissues. For m^5^C, we used atlas sites derived from bisulfite sequencing across 22 cell lines/tissues.

##### Sequence window construction

For each annotated modification site, we extracted a fixed-length sequence window of 41 nucleotides centered on the modified base (position 21). Windows extending beyond transcript boundaries were discarded. To ensure consistent tokenization across models, all sequences were represented in the {A,C,G,U} alphabet and mapped to {A,C,G,T} by converting U→T.

##### Positive sets

The m^6^A dataset contains 116,258 positive sites and the m^5^C dataset contains 134,623 positive sites (after preprocessing).

##### Negative sampling

Negatives were sampled from the same transcript as each positive site to control for transcript- and context-specific sequence composition. Specifically, for each m^6^A-positive adenosine, we randomly sampled unmodified adenosines from the same transcript that are not annotated as m^6^A in the atlas; analogously, for each m^5^C-positive cytidine, we sampled unmodified cytidines from the same transcript that are not annotated as m^5^C. We constructed multiple class-balance settings by sampling negatives to achieve positive:negative ratios of 1:1, 1:3, and 1:5. When multiple candidate negatives were available, sampling was performed uniformly at random without replacement; when insufficient candidates existed in a transcript, additional negatives were drawn from other transcripts with matching base identity to maintain the target ratio.

##### Evaluation

We trained and evaluated models under each class-balance setting using the same fine-tuning protocol. Because the task is highly imbalanced in realistic settings, we report area under the precision–recall curve (AUPRC) as the primary metric, together with F1/MCC where applicable (Supplementary Table S6).

#### RNA–RNA interaction data

We evaluate miRNA–target interaction prediction using the DeepMirTar benchmark [32]. DeepMirTar constructs positive miRNA–mRNA target-site pairs by integrating experimentally supported interactions including CLASH chimeric reads [36] and curated resources such as mirMark [37]. Negative pairs are generated by pairing/shuffling mature miRNAs against non-target sites, producing a balanced binary classification dataset (dataset statistics and splits follow [32]). DeepMirTar contains 13,860 positive and 13,860 negative miRNA–mRNA pairs; miRNAs are all shorter than 26 nt and the corresponding mRNA segments are shorter than 53 nt[32]. Following the RNAErnie’s setup, we report the accuracy, recall, precision, F1 and AUC. (see Supplementary Table S7)

#### mRNA stability/half-life data

To evaluate sequence determinants of stability/half-life, we include an mRNA stability regression task based on large-scale mRNA decay/stability measurements aggregated across vertebrate systems and cell types. In particular, we follow published compendia and predictive frameworks that integrate transcriptome-wide stability profiles across multiple species (human, mouse, frog and fish) and experimental contexts to model decay from sequence features [38, 39].The resulting dataset contains 41,123 sequences with lengths ranging from 30 to 1,497 nucleotides. To align with CodonBERT[21], we report Spearman’s rank correlation (*ρ*) (see Supplementary Table S8).

#### Translation efficiency (TE) and expression level (EL) data

For TE/EL prediction, we use matched RNA-seq and Ribo-seq measurements curated from public ribosome profiling studies accessible via GWIPS-viz [40, 41]. For each transcript, we extract a 100-nt sequence window immediately upstream of the annotated start codon (5^′^UTR-proximal region). Expression level (EL) targets are derived from RNA-seq as a proxy for mRNA abundance, and translation efficiency (TE) targets are derived from Ribo-seq as ribosome density normalized by EL (i.e., TE ∝ Ribo-seq*/*RNA-seq). Our final datasets included 14,410 sequences from HEK cells, 12,579 sequences from PC3 cells, and 1,257 sequences from muscle tissue. Model performance for both EL and TE is reported using Spearman’s rank correlation to enable robust comparison across datasets and cell types (Supplementary Table S9).

#### Mean ribosome loading (MRL) data

For MRL prediction, we use massively parallel polysome-profiling reporter assays that quantify mean ribosome loading as a sequence-determined proxy for translation [42]. In particular, we use the human 5^′^UTR library comprising 35,212 truncated endogenous human 5^′^UTRs together with 3,577 naturally occurring variants [42]. Unless otherwise stated, we follow the BEACON protocol for train/validation/test splits, and report both Pearson *R*^2^ and Spearman’s rank correlation (*ρ*) for performance comparison. (Supplementary Table S10).

#### RBP motif analysis

For each RBP, we ranked all test-set sites by the model-predicted binding probability and selected the top 1,000 high-confidence predicted binding sites and 1,000 high-confidence predicted non-binding sites. Specifically, predicted positives were required to have probability *>* 0.9, while predicted negatives were required to have probability *<* 0.1. For each site, we extracted a 101-nt sequence window centered on the peak midpoint (50 nt upstream and 50 nt downstream). We then performed de novo motif enrichment using MEME [43] on the predicted positives and negatives, and compared the resulting sequence logos to known RBP motif logos downloaded from the ATtRACT database[44].

## Supplementary Information

**Table S1.** RNAElectra architecture and optimization hyperparameters.

| Parameter | Generator | Discriminator (RNAElectra) |
| --- | --- | --- |
| Number of layers | 12 | 22 |
| Hidden size | 256 | 512 |
| Attention heads | 8 | 16 |
| Intermediate size | 1024 | 2048 |
| Position embeddings | Rotary | Rotary |
| Attention implementation | FlashAttention-2 | FlashAttention-2 |
| Total parameters | ~15M | ~93M |
| Masking ratio | 15% | N/A |
| Sequence length | 1024 tokens | 1024 tokens |
| Learning rate | $1 \times 10^{-4}$ | $1 \times 10^{-4}$ |
| Optimizer | AdamW | AdamW |
| AdamW $\beta$ | (0.9, 0.999) | (0.9, 0.999) |
| AdamW $\epsilon$ | $1 \times 10^{-8}$ | $1 \times 10^{-8}$ |
| Weight decay | $3 \times 10^{-7}$ | $3 \times 10^{-7}$ |

**Table S2.** RNAElectra Pretraining Data Summary.

| Item | Value | Notes |
| --- | --- | --- |
| Data source | RNAcentral (Active) | Integrated from Ensembl, Rfam, RefSeq |
| Input sequences | 45,316,669 | Raw downloaded sequences |
| Valid sequences | 44,613,362 | No N and length $\leq 1024$ nt |
| Filtered out | 703,307 (1.55%) | Removed due to N |
| Train / validation split | 90% / 10% | Random split |
| Min / max length | 10 / 1024 nt | Length measured in nucleotides |
| Mean length | 445.37 nt | Over valid sequences |

**Table S3.** Common training configuration used for most BEACON tasks.

| config | value |
| --- | --- |
| optimizer | AdamW |
| optimizer $\epsilon$ | $1 \times 10^{-8}$ |
| optimizer momentum | $\beta_1=0.9, \beta_2=0.999$ |
| weight decay | 0.01 |
| learning-rate schedule | cosine decay |
| random seeds | [30, 42, 2026] |
| learning rate | $[1 \times 10^{-5} - 5 \times 10^{-3}]$ |
| warmup steps | 50 |
| epochs | 30 |
| batch size | 32 |
| gradient accumulation | 1 |
| dtype | float16 |

**Table S4.** Performance comparison of RNA secondary structure prediction methods on the ArchiveII600 dataset.

| Methods | Precision | Recall | F1 |
| --- | --- | --- | --- |
| RNAstructure | 0.563 | 0.615 | 0.585 |
| RNAsoft | 0.665 | 0.594 | 0.622 |
| RNAfold | 0.565 | 0.627 | 0.592 |
| MXfold2 | 0.788 | 0.760 | 0.768 |
| Mfold | 0.428 | 0.383 | 0.401 |
| LinearFold | 0.641 | 0.617 | 0.621 |
| Eternafold | 0.667 | 0.622 | 0.636 |
| E2Efold | 0.738 | 0.665 | 0.690 |
| Contrafold | 0.607 | 0.679 | 0.638 |
| Contextfold | 0.873 | 0.821 | 0.842 |
| RNABERT | 0.634 | 0.649 | 0.641 |
| RNA-MSM | 0.664 | 0.648 | 0.656 |
| RNA-FM | 0.752 | 0.737 | 0.744 |
| Ernie-base | 0.875 | 0.839 | 0.851 |
| RNAErnie<sup>-</sup> | 0.855 | 0.844 | 0.846 |
| RNAErnie<sup>+</sup> | 0.848 | 0.854 | 0.848 |
| RNAErnie | 0.884 | 0.869 | 0.873 |
| RNAErnie<sup>+</sup> | 0.886 | 0.870 | 0.875 |
| RNAElectra | 0.960 | 0.918 | 0.933 |

**Table S5.** Comparison of AUROC performance across different methods under Neg-1 and Neg-2 setups. RNAElectra achieves the best performance among all sequence-only language models.

| Method | Target | Additional Input | Neg-1 | Neg-2 | $\Delta$ |
| --- | --- | --- | --- | --- | --- |
| *Sequence-only* |  |  |  |  |  |
| BERT-RBP | Binary | Sequence | 0.8927 | 0.8112 | -0.0815 |
| Pysster-101 | Binary | Sequence | 0.9003 | 0.8311 | -0.0692 |
| Pysster | Binary | Sequence | 0.9190 | 0.8497 | -0.0693 |
| DeepRAM | Binary | Sequence | 0.8697 | 0.8029 | -0.0668 |
| DeepCLIP | Binary | Sequence | 0.8733 | 0.8081 | -0.0652 |
| GraphProt | Binary | Sequence | 0.8211 | 0.7756 | -0.0455 |
| MultiRBP | Multitask | Sequence | NA | 0.7937 | NA |
| RNAElectra | Binary | Sequence | 0.9068 | 0.8570 | -0.0498 |
| *Sequence + Additional Features* |  |  |  |  |  |
| PRISMNet | Binary | icSHAPE | 0.9015 | 0.8255 | -0.0760 |
| iDeepS | Binary | Predicted structure | 0.8932 | 0.8285 | -0.0647 |
| RNAProt-Extended | Binary | Conservation, region, struct | 0.9419 | 0.8569 | -0.0850 |
| DeepRiPe | Multitask | Region features | 0.9103 | 0.8774 | -0.0329 |
| Multi-resBind | Multitask | Region features | 0.9174 | 0.8825 | -0.0349 |

**Table S6.** Epitranscriptomic modification site prediction (PRAUC). Best and second-best per row are marked by and, respectively.

| Task | Pos:Neg | ErnieRna | RNAErnie | RNAFM | RiNALMo | RNAElectra |
| --- | --- | --- | --- | --- | --- | --- |
| m<sup>5</sup>C | 1:1 | 0.8953 | 0.8658 | 0.8847 | 0.8810 | 0.9156 |
|  | 1:3 | 0.7744 | 0.7019 | 0.7448 | 0.7094 | 0.8442 |
|  | 1:5 | 0.6562 | 0.5766 | 0.6390 | 0.6814 | 0.8013 |
| m<sup>6</sup>A | 1:1 | 0.8598 | 0.8394 | 0.8607 | 0.8637 | 0.8582 |
|  | 1:3 | 0.7161 | 0.6823 | 0.7183 | 0.7169 | 0.7096 |
|  | 1:5 | 0.6338 | 0.5926 | 0.6362 | 0.6391 | 0.6342 |

**Table S7.** Performance of RNAElectra on RNA–RNA interaction prediction using the DeepMirTar dataset.

| Method | Accuracy | Recall | Precision | F1 | AUC |
| --- | --- | --- | --- | --- | --- |
| Miranda | 0.6592 | 0.6522 | 0.6662 | 0.6591 | 0.6874 |
| RNAhybrid | 0.6988 | 0.6446 | 0.7535 | 0.6948 | 0.7585 |
| PITA | 0.4981 | 0.5872 | 0.4082 | 0.4816 | – |
| TargetScan v.7.0 | 0.5801 | 0.6023 | 0.5922 | 0.5972 | 0.6725 |
| TarPmiR | 0.7446 | 0.7368 | 0.7656 | 0.7509 | 0.8021 |
| DeepMirTar | 0.9348 | 0.9235 | 0.9479 | 0.9245 | 0.9793 |
| RNABERT | 0.8375 | 0.8372 | 0.8378 | 0.8373 | 0.9160 |
| RNA-MSM | 0.8205 | 0.8203 | 0.8207 | 0.8204 | 0.9048 |
| RNA-FM | 0.9208 | 0.9208 | 0.9208 | 0.9208 | 0.9741 |
| Ernie-base | 0.9262 | 0.9257 | 0.9371 | 0.9314 | 0.9674 |
| RNAErnie<sup>–</sup> | 0.9537 | 0.9547 | 0.9533 | 0.9540 | 0.9801 |
| RNAErnie<sup>+</sup> | 0.9524 | 0.9539 | 0.9514 | 0.9526 | 0.9811 |
| RNAErnie | 0.9570 | 0.9576 | 0.9571 | 0.9573 | 0.9876 |
| RNAElectra | 0.9650 | 0.9589 | 0.9724 | 0.9656 | 0.9938 |

**Table S8.**
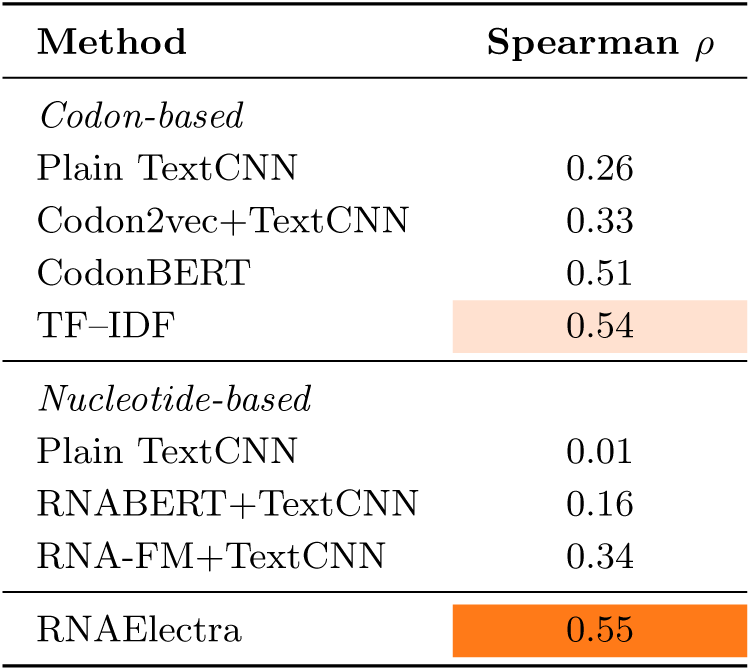
mRNA stability prediction performance measured by Spearman correlation (*ρ*).

**Table S9.** Performance comparison on Translation Efficiency (TE) and Expression Level (EL) prediction using Spearman correlation across Muscle, PC3, and HEK293T datasets.

| Method | TE (Spearman $\rho$) | | | EL (Spearman $\rho$) | | |
| --- | --- | --- | --- | --- | --- | --- |
|  | Muscle | PC3 | HEK | Muscle | PC3 | HEK |
| RNABERT_MLP | 0.32 | 0.32 | 0.23 | 0.20 | 0.20 | 0.22 |
| RNAFM_MLP | 0.30 | 0.54 | 0.50 | 0.30 | 0.48 | 0.47 |
| Kipoi | 0.31 | 0.34 | 0.35 | 0.28 | 0.33 | 0.37 |
| Optimus | 0.41 | 0.38 | 0.36 | 0.15 | 0.19 | 0.18 |
| MTtrans | 0.50 | 0.54 | 0.50 | — | — | — |
| Cao-RF | 0.63 | 0.63 | 0.55 | 0.64 | 0.59 | 0.57 |
| UTR-LM | 0.67 | 0.65 | 0.60 | 0.66 | 0.63 | 0.65 |
| RNAElectra | 0.68 | 0.69 | 0.63 | 0.65 | 0.64 | 0.65 |

**Table S10.**
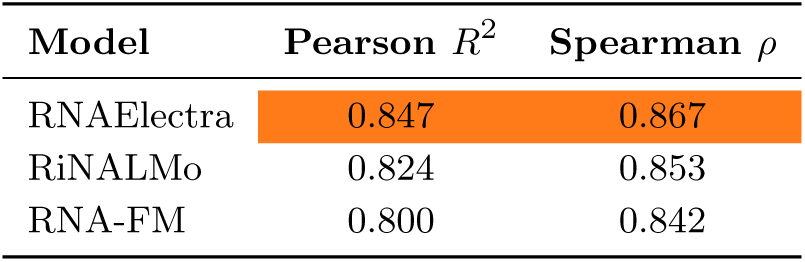
mRNA translation efficiency (MRL) prediction performance.

## Data availability

The datasets used supporting the conclusions of this article are referenced within the article. RNAcentral sequences used for pretraining are publicly available from the RNAcentral FTP server: https://ftp.ebi.ac.uk/pub/databases/RNAcentral/current_release/sequences/rnacentral_active.fasta.gz. The BEACON benchmark datasets and task definitions are available at https://github.com/terry-r123/RNABenchmark. Additional downstream datasets used in this work are available from the following sources: RNAStrAlign (https://github.com/mxfold/mxfold2/releases/tag/v0.1.0), Archivell (https://drive.google.com/drive/folders/1Sq7MVgFOshGPlumRE_hpNXadvhJKaryi?usp=sharing), bpRNA-1m (https://bprna.cgrb.oregonstate.edu/download.php#bpRNA), Deep-MirTar (https://github.com/tjgu/miTAR/tree/master/scripts_data_models), RBP Benchmark (https://github.com/mhorlacher/Benchmark-RBP), m^5^C-Atlas (http://180.208.58.19/m5c-atlas/), m^6^A-Atlas v2.0 (https://rnamd.org/m6a/), mRNA stability data (https://github.com/santiago1234/iCodon), translation efficiency data (https://drive.google.com/drive/folders/1oGGgQ33cbx340vXsH_Ds_Py6Ad0TslLD), and ATtRACT (https://attract.cnic.es/index). The RNAElectra pretrained model weights are available via Hugging Face (`FreakingPotato/RNAElectra`).

## Acknowledgements

We acknowledge the National Computational Infrastructure (NCI), Australia, for providing computational resources and support.

## Declarations

### Ethics approval and consent to participate

Not applicable.

